# SeedVicious: analysis of microRNA target and near-target sites

**DOI:** 10.1101/124529

**Authors:** Antonio Marco

**Author notes:** To whom correspondence should be addressed School of Biological Sciences University of Essex Wivenhoe Park, Colchester CO4 3SQ United Kingdom, Contact.

## Abstract

Here I describe seedVicious, a versatile microRNA target site prediction software that can be easily fitted into annotation pipelines and run over custom datasets. SeedVicious finds microRNA canonical sites plus other, less efficient, target sites. The program also detects near-target sites, which have one nucleotide different from a canonical site. Near-target sites are important to study population variation in microRNA regulation. Here I show that near-target sites can also be functional sites. Among other features, seedVicious can also compute evolutionary gains/losses of target sites using maximum parsimony. SeedVicious does not aim to outperform but to complement existing microRNA prediction tools. For instance, the precision of TargetScan is doubled (from 11% to ~22%) when we filter predictions by the distance between target sites using our program. The software is written in Perl and runs on 64-bit Unix computers (Linux and MacOS X). Users can also try the program in a dedicated web-server by uploading custom data, or browsing pre-computed predictions. SeedVicious and its associated web-server and database (SeedBank) are distributed under the GPL/GNU license.

## INTRODUCTION

Animal microRNAs target gene transcripts by partial sequence complementarity (1). There are multiple microRNA target prediction tools that use different strategies. Most prediction programs look at the existence of seed sequences, six-nucleotide-long sequences in the transcripts that are complementary to nucleotides 2 to 7 in the microRNA (1). Depending on additional nucleotide pairings these sites can be canonical or marginal. Many programs exploit additional features, mainly evolutionary conservation and RNA folding features (2, 3). Although these programs have an increased accuracy, they lose power as many real targets remain undetected. Often, microRNA target predictions are available only on selected datasets, and stand-alone programs are not available for use on custom data. Here I describe a new microRNA target prediction software that does not aim to replace but to complement the existing tool-kit, and allow the high-throughput analysis of custom microRNA/transcript data as well as the exploration of additional features not covered by other programs.

The first population genetics studies on microRNA targets sites already described selective pressures on seed sequences (4, 5). In a recent study I showed that, in *Drosophila* populations, there is selection against microRNA target sites (6). To study selection on non-target sites that may become targets, I defined ‘one-mutant neighbors’ as six-nucleotide-long sequences that have one nucleotide different to a seed sequence. Here I define a broader term: near-target sites (Figure 1), which have one nucleotide different to a putative microRNA target site, and are not targets themselves. Detection of near-target sites are important for population genetics and evolutionary studies, and will allow us to explore the selective pressures on gene transcripts. Additionally, reference genome sequences, in which microRNA target sites are usually predicted, do not capture the true diversity of a species and may miss *bona fide* target sites. Scanning for near-target sites may reveal targets that would be otherwise ignored. SeedVicious allows the exploration of near-target sites.

**Figure 1.**
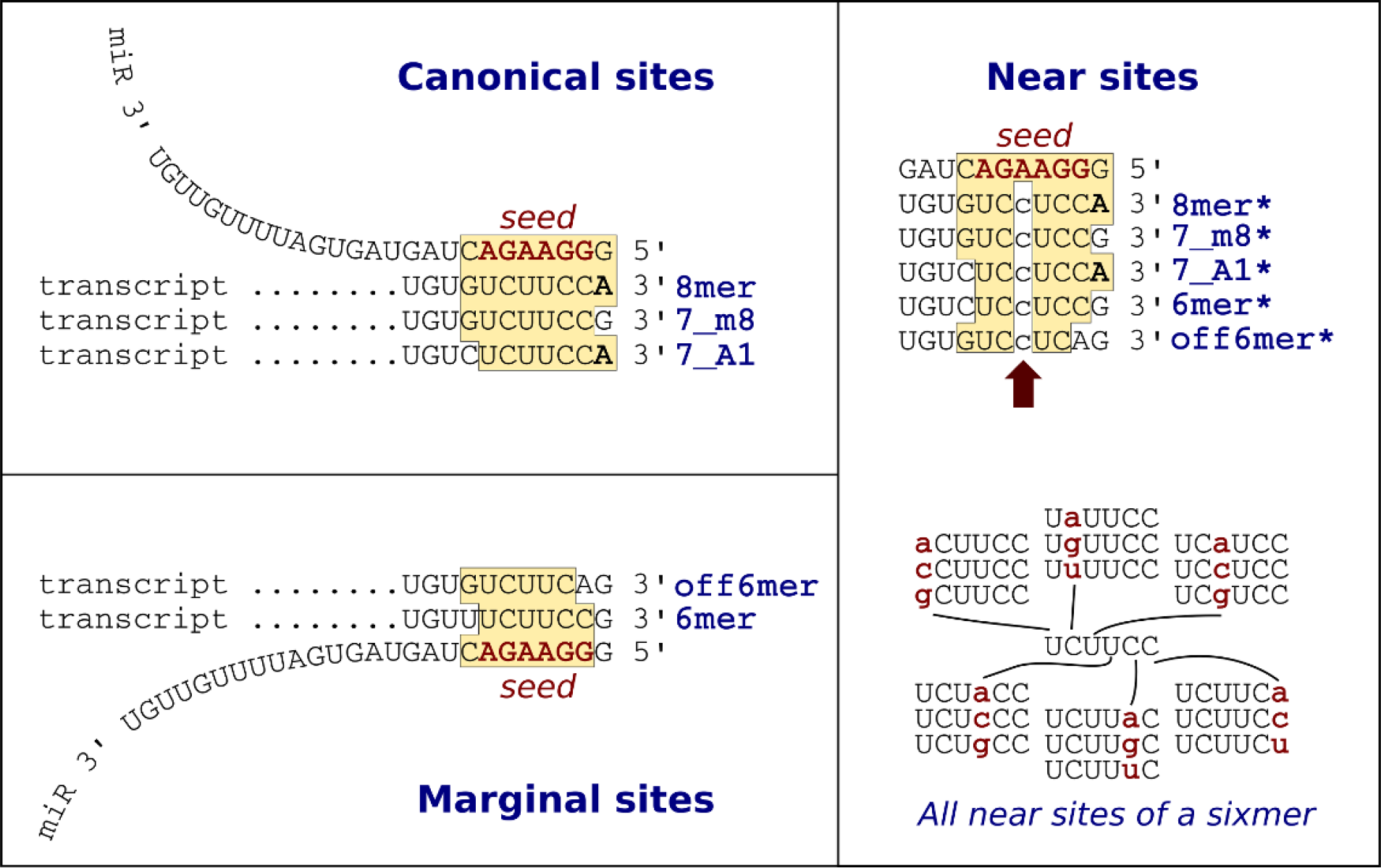
MicroRNA target and near-target sites. Canonical and marginal sites are described in Bartel (2009). Near-target sites can be defined for all type of sites. The figure also shows that each seed (sixmer) 18 possible near-target sites as defined by Marco (2015).

## METHODS AND IMPLEMENTATION

SeedVicious initially scans custom sequences to detect canonical microRNA target sites as described in (1). Optionally, marginal sites can be reported (Figure 1). This strategy produces a large number of false positives, and other prediction tools use evolutionary conservation to narrow down the number of potential target sites (2, 7). However, evolutionary conservation is not necessarily associated with functionality of the target site (8). SeedVicious allows an alternative (and complementary) way of finding putative target sites. The complete list of canonical (and marginal) sites can be ranked according to selected features. First, seedVicious computes the free energy of the microRNA:transcript duplex using the RNAeval program in the Vienna Package (9). The lower the energy, the more stable is the RNA:RNA pair, which can be interpreted as a more efficient target site. Second, seedVicious can also report the minimum distance between pairs of target sites for the same microRNA.

SeedVicious permits the inference of gains and losses of microRNA target sites by first predicting individual target sites at all transcripts in a given alignment, and then fitting a maximum parsimony (MP) model to a tree provided by the user, following Dollo’s criteria (10). The MP reconstruction of ancestral states is computed with the ‘dollop’ program from the Phylip package (11). We first used this method to study the evolution of post-transcriptional regulation in a gene family in Drosophila (12). A number of additional analyses can be performed with seedVicious. It can compare different 3’UTRs and detect either common microRNAs targeting pairs of transcripts or common target sites in an alignment. A major feature of seedVicious is the detection of near-target sites which are detected for both canonical and marginal target sites.

The main program can be run from the command line using Perl 5 or above, and the required modules and external binary files (compiled in a 64-bit Unix computer) are included in the distributed version. Input sequence files should be in FASTA format, and tree files in NEWICK format. Input files can be compressed in Gzip format. A fully referenced User Guide is available form the package or as Supplementary Information to this paper, and it includes different protocols.

For the analysis presented below, all microRNAs were retrieved from miRBase release 21 (13), and all transcript sequences from ENSEMBL Genes 89 via Biomart (14). TargetScan targets were retrieved from targetscan.org, release 7.1 (2). Predictions were compared to the experimentally validated targets in miRTarBase 6.0 (15), for both weak and strong interactions. Precision was calculated as the ratio TP/(TP+FP), where TP (True Positives) was the number of predicted targets that were experimentally validated, and FP (False Positives) was the number of predicted targets not validated in miRTarBase.

## RESULTS AND DISCUSSION

I describe in this section a few examples that illustrate different ways of analyzing target sites with seedVicious. First, I computed with seedVicious the minimum distance between canonical target sites for all human microRNAs in 3’UTRs (see Methods and Implementation). This distance can be used as a filtering criteria for potential targets, based on a recent work that suggests that neighboring sites cooperate during lateral diffusion of the Ago-miRNA complex during target recognition (16). Overall, the minimum distance between target sites for microRNA/transcript pairs with experimentally validated targets is smaller than for non-validated targets (Figure 2A). This indicates that the closest two target sites are in a transcript, the more likely that the microRNA will regulate the targeted transcript. As a matter of fact, two contiguous target sites (6 nt distance) significantly increases the chances of work as a regulatory sequence (Figure 2B).

**Figure 2.**
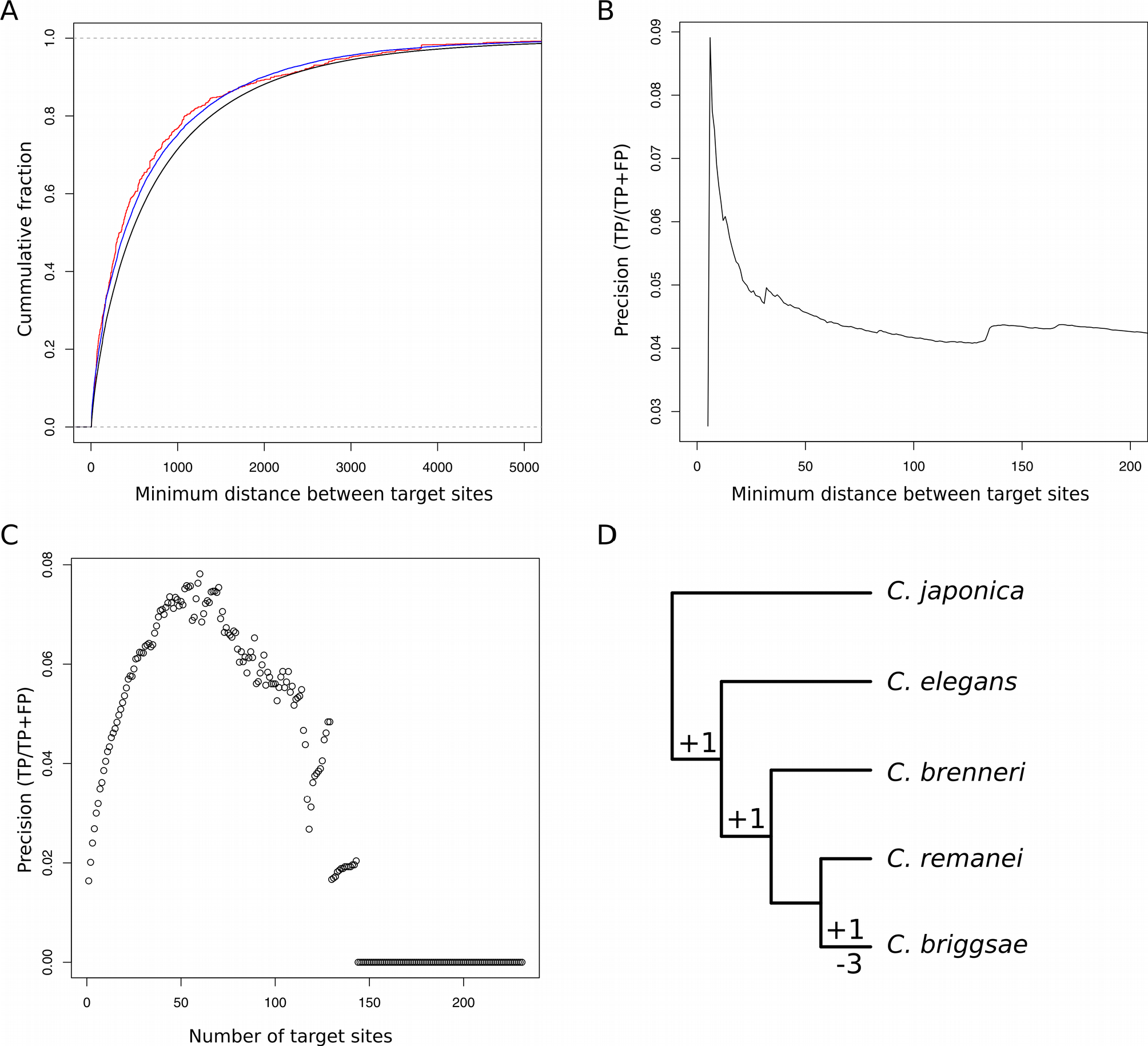
Multiple uses of seedVicious. A) Cummulative distribution of the number of transcripts with at least tow target sites with respect to the minimum distance between pairs of targets. The graph compares the distribution of all transcript (black), transcript/microRNA pairs with experimentally validated interactions (blue) and those with strongly validated interactions (red) according to miRTarBase. B) Precision of microRNA target prediction for different cut-offs of minimum distance between pairs of targets. The peak corresponds to a distance of 6, that is, two contiguous target sites. C) Precision of microRNA target prediction for multiple targets sites for the same microRNA. D) Evolutionary turnover of microRNA canonical target sites for let-7 in lin-14 in roundworm species using maximum parsimony.

The precision of a target prediction is measured as the fraction of true positives with respect to the total number of predictions. By using a compilation of experimentally validated microRNA/transcript interactions (see Methods and Implementation) I estimated the precision of target prediction based on the minimum distance between canonical sites. If we predict as microRNA targets transcripts with at least 2 canonical sites, the precision of the prediction is 0.041. This value significantly increases (to 0.087) if we only consider pairs of canonical sites next to each other (distance of 6 nucleotides). Precision values in microRNA target prediction are generally low, in particular when one uses a small dataset of validated targets (as in this case), but it is useful to compare the predictive power of different strategies. Using this same dataset, the precision of TargetScan (using conserved targets only) is 0.11. Strikingly, if we combine TargetScan and pairs of targets at 6 nucleotides in distance, the precision goes up to 0.21. In conclusion, combining some of the features of seedVicious to other programs can be use to increase the precision in microRNA target prediction.

In previous studies, the occurrence of two or more seed sequences in transcripts has been associated to a higher chance of predicting a *bona fide* targeting interaction (17). Here I compared the proportion of predicted targets that are validated by experiments (precision) and the number of canonical target sites. The graph in Figure 2C shows that, indeed, as the number of target sites increase, the proportion of validated targets also increases, although there is a global maximum around 50. It will be interesting to explore whether distinct microRNAs have a different optimum number of target sites for an efficient repression of the target transcript.

The analysis of targets and near-targets is of particular interest to analyse populations with segregating alleles in target sites (4, 6). For instance, the longest transcript from tumour-suppressor gene *p53* have no target sites (neither canonical nor marginal) for miR-7-5p, a well known cancer-related microRNA (18). However, it contains three near-target sites. That is, there are three potential nucleotide substitutions that will create a novel target site for this particular microRNA. Indeed, miR-7 promotes tumour growth via a mechanisms that includes p53 (19). Somatic mutations in the 3’UTR of p53 leading to new miR-7-5p target sites in cancer patients is then plausible. Current work in our lab reveals that selective pressures against this type of substitutions in cancer-associated genes is prevalent (Hatlen and Marco, unpublished results).

The transcript from the gene *lin-14* in *Caenorhabditis elegans* has seven targets sites for the microRNA *lin-4* (20). However, other prediction programs detect only three (TargetScan) or none, probably due to the stringent criteria. Using seedVicious I scanned the lin-14 3’ UTR (21) for canonical target and canonical near-target sites. Three canonical sites were detected, the same that are reported in TargetScan. Additionally, five near-target sites were reported, four of them corresponding to the other four sites originally described. One of the near-target sites was not previously described and may be a good candidate to further explore. The detailed analysis as well as the input files are available with the package and fully described in the User Guide (Supplementary File 1). This example illustrates the potential of studying near-target sites, not only in evolutionary studies, but in the detection of potential functional targets. Further examples are described in the software manual.

The canonical target sites for lin-4-5p in lin-14 are highly conserved. Actually, in five species of worm studies all sites remain the same. However, let-7-5p, which also targets lin-14, has a more dynamic evolution. Using seedVicious I predicted the canonical target sites in five worm species and to reconstruct the gains and losses using maximum parsimony. Results are shown in Figure 2C, where we observe that there have been 4 target sites gained among the studied species whilst *Caenorhabditis briggsae* lost its three ancestral target sites.

In conclusion, seedVicious is a valuable addition to the toolkit of microRNA biologists. It works on custom datasets and can be easily fitted into annotation pipelines. It provides a range of analysis that can be combined with other programs to generate robust microRNA target predictions. The analysis of near-target sites is also useful to study population dynamics at target sites, and the inbuilt maximum parsimony functionality is valuable to study the evolution of microRNA-mediated regulation at a large scale.

## AVAILABILITY

The program is available for download at http://seedvicious.essex.ac.uk/download.html. A web version is available, which allows users to run the program using our server. Precomputed targets for selected species can also be browsed from our SeedBank database. The web server and the database are accessed via http://seedvicious.essex.ac.uk

## ACKNOWLEDGEMENTS

This software would not have been made publicly available without the encouragement and support of my colleagues, in particular Mohab Helmy and Sam Griffiths-Jones. I am also grateful to M. Helmy and Andrea Hatlen for discussion, to Roman Cheplyaka for feedback on the program, and to Stuart Newman for his invaluable help setting up the web-server.

